# The flavin monooxygenase Bs3 triggers cell death in plants, impairs growth in yeast and produces H_2_O_2_ *in vitro*

**DOI:** 10.1101/2021.02.03.429511

**Authors:** Christina Krönauer, Thomas Lahaye

## Abstract

The pepper resistance gene *Bs3* triggers a hypersensitive response (HR) upon transcriptional activation by the corresponding transcription activator-like effector AvrBs3 from the bacterial pathogen *Xanthomonas*. Bs3 is homologous to flavin monooxygenases (FMOs), an enzyme class that has NADPH oxidase activity and can produce H_2_O_2_, a hallmark metabolite in plant immune reactions. Histochemical staining of infected pepper leaves and a translational fusion of Bs3 to the redox reporter roGFP2 both indicated that the Bs3-dependent HR induces a local increase in H_2_O_2_ levels *in planta*. Moreover, our *in vitro* studies with recombinant Bs3 protein confirmed its NADPH oxidase activity. To test if the NADPH oxidation of Bs3 induces HR, we adapted previous studies which have uncovered mutations in fungal FMOs that result in higher NADPH oxidase activity. We replicated one of these mutations and demonstrated that the generated recombinant Bs3_S211A_ protein has twofold higher NADPH oxidase activity than wildtype Bs3 *in vitro*. Translational fusions to roGFP2 showed that Bs3_S211A_ also had increased NADPH oxidase activity *in planta*. Interestingly, while the mutant derivative Bs3_S211A_ had an increase in NADPH oxidase capacity, it did not trigger HR *in planta*. Ultimately, this reveals that Bs3 produces H_2_O_2_ *in planta*, but that the H_2_O_2_ produced by Bs3 on its own is not sufficient to trigger HR. We also demonstrated that expression of Bs3 not only triggered HR in plants, but also inhibited proliferation of yeast, which lends this model system to be utilized for the genetic dissection of Bs3 function in future studies.

**One sentence summary:** The executor-type resistance protein Bs3 from pepper (*Capsicum annuum*) acts as an NADPH oxidase but reactive oxygen species produced by Bs3 are not sufficient to trigger plant cell death

## INTRODUCTION

Programmed cell death (PCD) is a key component of the plant immune system and provides protection against biotrophic microbial pathogens. Execution of pathogen-triggered plant cell death, often referred to as the hypersensitive response (HR), is generally controlled by two distinct immune receptor classes, membrane-resident pattern recognition receptors (PRRs) and intracellular nucleotide-binding domain leucine-rich repeat (NLR) proteins, which are the most abundant type of plant resistance (*R*) proteins (Zhou and Zhang, 2020). Upon recognition of pathogen structures or pathogen-induced changes in the host cell, these receptors initiate a number of cellular events, like calcium influx, burst of reactive oxygen species (ROS), and accumulation of salicylic acid (SA) that are assumed to serve as signal molecules that eventually trigger HR (Noctor and Mhamdi, 2017). It is unclear, however, how the activation of plant immune receptors eventually translates into a cell death reaction.

We study the *R* gene *Bs3* from pepper (*Capsicum annuum*) which mediates recognition of the *Xanthomonas* transcription activator like effector (TALE) protein, AvrBs3 (Römer et al., 2007). TALEs are one class of bacterial effectors that, upon injection into host cells, translocate to the plant nucleus where they bind to an approximately 20 base pair long effector binding element (EBE) and transcriptionally activate the downstream host susceptibility (*S*) gene to promote disease (Boch et al., 2014). Some genotypes of susceptible plant species have evolved a functionally unique class of *R* genes, the so-called executor *R* genes. Such *R* genes contain EBEs in their promoters that mediate interaction and transcriptional activation by TALEs. TALEs are perceived by EBEs present in the *R* gene promoters, and the downstream TALE-induced expression of the executor R protein results in the execution of a plant immune reaction (Timilsina et al., 2020). Hence, executor-type *R* genes have distinct functional modules for effector recognition (*R*-gene promoter) and orchestration of the immune response (executor R protein). This is in striking contrast to NLR type R proteins, which are known to mediate both, detection of effectors and induction of an immune reaction. At the mechanistic level, there are resemblances between executor type R proteins and activated NLRs. However, whether or not executors employ canonical immune pathways that are utilized by activated NLRs needs to be elucidated.

Five executor-type *R* genes have been cloned so far. With exception of the rice executor R proteins Xa10 and Xa23 that share about 50% sequence identity to each other, executor R proteins show neither sequence relatedness to each other nor homology to any other NLR-type or PRR-type immune receptor. Within the class of executor R proteins, Bs3 is exceptional as it is the only one that shares homology to a protein class of known function. More specifically, Bs3 shows homology to flavin-containing monooxygenases (FMOs), a class of enzymes that uses molecular oxygen (O_2_) for oxygenation of metabolites (Krueger and Williams, 2005). In plants, FMOs are a large and diverse group, but only a few members that have been found to function in hormone production or pathogen defense have been characterized so far (Dai et al., 2012; Hartmann et al., 2018; Thodberg et al., 2020). In contrast to other plant FMOs that have a function in immune-signaling or chemical defense, Bs3 is unique as it is the only FMO known to trigger cell death. Bs3 is most closely related to YUCCA proteins, a plant-specific FMO subgroup that catalyzes the final step in tryptophan-dependent auxin (Indole-3-Acetic Acid; IAA) biosynthesis (Dai et al., 2013). The most prominent difference of Bs3 and YUCCAs is a stretch of ~70 amino acids, that is conserved across all YUCCAs but absent from Bs3 (Krönauer et al., 2019). Studies of mammalian FMOs established structure-function relationships with several motifs being conserved also in plant YUCCAs (Thodberg and Jakobsen Neilson, 2020), however, no function has been assigned to this ~70 residue stretch absent in Bs3 and ultimately its functional relevance cannot be foreseen.

Given that Bs3 is an FMO, we took advantage of the well characterized FMO enzymatic cycle to gain insight into how Bs3 could potentially trigger cell death in plants. We focused on the reduction of the bound FAD cofactor in FMOs by NADPH and binding of molecular oxygen that results in a C4a-(hydro)peroxyflavin (C4a) intermediate, which is ready to oxygenate suitable substrates. If no metabolic substrate is available, the FMO C4a intermediate breaks down without substrate oxygenation, a process referred to as the uncoupled reaction, where reduction equivalents are released as H_2_O_2_ (Alfieri et al., 2008). While it is unclear if the production of H_2_O_2_ by FMOs is an undesired waste of NADPH, or if it potentially serves a biological function, the role of H_2_O_2_ as a signaling molecule in plant immune reactions is well established (Siddens et al., 2014; Waszczak et al., 2018). We speculated that NADPH oxidase activity of Bs3 and subsequent release of H_2_O_2_ could play a role in induction of HR.

In this study, we tested if synthesis of H_2_O_2_ by Bs3 is the metabolic trigger that initiates Bs3-mediated HR. Staining of *Xanthomonas* infected pepper leaves with 3,3-diaminobenzidine tetrahydrochloride (DAB) indicates that the Bs3-dependent HR correlates with the accumulation of reactive oxygen species (ROS). *In vitro* studies of recombinant Bs3 protein and *in planta* studies with the redox-sensitive roGFP2 suggest that Bs3 has NADPH oxidase activity, however, a Bs3-mutant derivative with elevated NADPH oxidase activity failed to trigger HR *in planta*. This observation is not consistent with a model where ROS produced by Bs3 is the metabolic trigger of Bs3-dependent HR. We also show that Bs3 not only triggers HR in plant cells but also inhibits proliferation of yeast, a model system that might be a promising experimental platform to genetically dissect Bs3-dependent cell death in future studies.

## RESULTS

### Execution of Bs3 HR correlates with accumulation of H_2_O_2_ in planta

Our previous studies have shown that despite its high similarity to auxin-producing YUCCA proteins, pepper Bs3 does not induce accumulation of auxin (Krönauer et al., 2019). Thus, other enzymatic features of YUCCA proteins were inspected with respect to their potential to possibly trigger HR. In general, FMOs have NADPH oxidase activity and can, depending on the stability of their C4a intermediate (Chaiyen et al., 2012), synthesize significant amounts of H_2_O_2_, a key metabolite of plant immune reactions (Mignolet-Spruyt et al., 2016; Mittler, 2017). We accordingly hypothesized that Bs3 is an FMO with high NADPH oxidase activity, which triggers HR by production of H_2_O_2_. To clarify if Bs3 expression indeed correlates with increased H_2_O_2_ levels *in planta*, we inoculated leaves of the pepper genotype Early Calwonder 123 (ECW123; [Stall R.E.; Jones et al., 2002]), that contains the bacterial spot (Bs) plant *R* genes *Bs2* and *Bs3* with a *Xanthomonas euvesicatoria* (*Xeu*) strain that lacks the *avrBs2* and *avrBs3* genes (82-8 universally susceptible [uns]) or isogenic transconjugants containing either *avrBs2* or *avrBs3*. To visualize possible accumulation of H_2_O_2_ within infected tissue, we performed histochemical staining of pepper leaves using 3,3-diaminobenzidine tetrahydrochloride (DAB) as described previously (Thordal-Christensen et al., 1997). Brown staining, indicative of accumulation of H_2_O_2_, was observed in leaf tissue infected with *Xeu* strains containing either AvrBs2 or AvrBs3, but not in leaf tissue infected with the *Xeu* strain 82-8 uns that lacks AvrBs2 and AvrBs3 (Fig. 1A). This shows that the execution of an HR by the NLR protein Bs2 or the executor R protein Bs3 in pepper leaves both correlate with a local increase of H_2_O_2_ levels. This observation is consistent with previous studies in *Nicotiana benthamiana* leaves where *Agrobacterium tumefaciens* mediated delivery (agroinfiltration) of a *35S* promoter-driven *Bs3* T-DNA caused ROS accumulation in inoculated leaf tissue after two days (Krönauer et al. 2019, Fig. S1). Our studies in pepper and *N. benthamiana* are consistent with a model in which Bs3 triggers HR via production of H_2_O_2_. However, whether H_2_O_2_ is directly produced by Bs3, or if a Bs3-dependent immune pathway eventually results in expression or activation of proteins that produce H_2_O_2_ cannot be determined from the conducted *in planta* assays.

**Figure 1:**
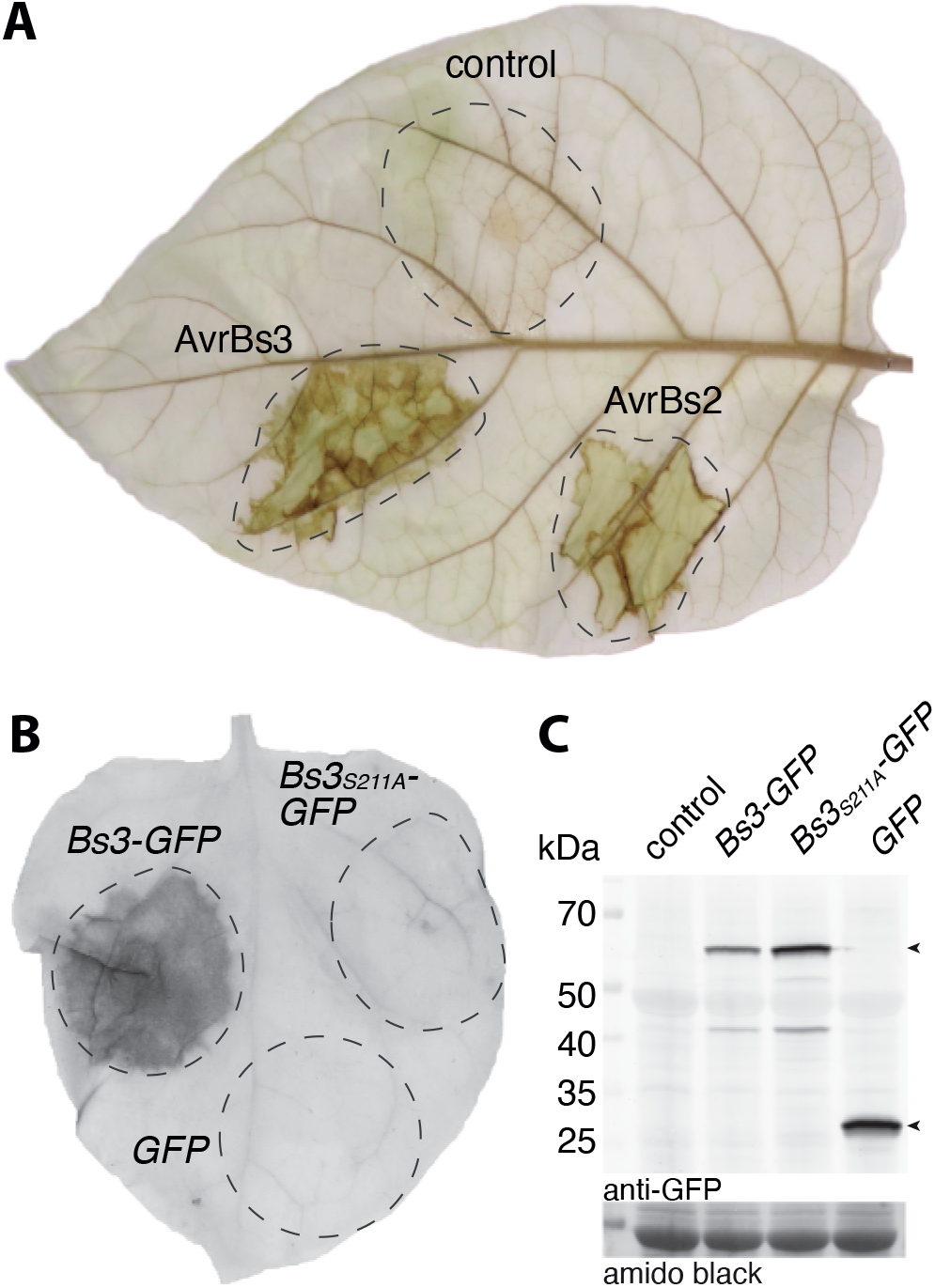
The Bs3-dependent HR correlates with H_2_O_2_ accumulation. A) Leaves of the *Capsicum annuum* genotype ECW123, which contains the bacterial spot (*Bs*) *R* genes *Bs2* and *Bs3* were infiltrated with the *Xanthomonas euvesicatoria* strain 82-8 uns (control) or corresponding transformants delivering the effector proteins AvrBs2 or AvrBs3. 30 hours post inoculation (hpi) the inoculated leaf was stained with 3,3-diaminobenzidine tetrahydrochloride (DAB) solution and cleared with ethanol. Dashed lines indicate inoculated leaf sections. B) *35S* promoter driven T-DNA constructs encoding Bs3-GFP, Bs3_S211A_-GFP or GFP were delivered into *N. benthamiana* leaves via *Agrobacterium*-mediated transient transformation. Four days post inoculation (dpi), the leaf was harvested and cleared in ethanol. The HR is visible as a dark area. C) *In planta* expression of T-DNA-encoded proteins was validated via an anti GFP immunoblot. The expected size for Bs3-GFP and GFP is indicated by arrows. Amido black staining was used to visualize total protein load.

### The NADPH binding site mutant Bs3_S211A_ does not trigger HR in planta

FMO function generally depends on transient binding of the NADPH cofactor, and mutations in codons that translate into conserved glycine residues within the NADPH binding site of FMOs have been shown to cause loss of enzyme activity (Lawton and Philpot, 1993; Hou et al., 2011). Mutational studies also uncovered specific mutation types within the NADPH binding site of FMOs that lead to derivatives producing increased amounts of H_2_O_2_ relative to the corresponding wild-type protein. For example, a serine to alanine mutation within the conserved NADPH binding site (G×G×**S**G ⟶ G×G×**A**G) of the *Aspergillus fumigatus* FMO SidA, results in a derivative with increased NADPH oxidase activity (Shirey et al., 2013). Inspired by these studies, we aimed to echo such mutations in the context of the pepper Bs3 protein to generate a mutant derivative with increased NADPH oxidase activity that would possibly induce a faster Bs3 HR. To do so, we mutated the triplet encoding the Bs3 serine at residue 211 to an alanine codon, to create Bs3_S211A_, a Bs3 derivative which would conceivably produce increased H_2_O_2_ levels. To test whether or not Bs3_S211A_ triggers HR *in planta*, we agroinfiltrated *35S* promoter driven *Bs3-GFP* or *Bs3*_*S211A*_*-GFP* into *N. benthamiana* leaves. These agroinfiltration assays showed that Bs3, but not Bs3_S211A_ triggered HR in *N. benthamiana* leaves (Fig. 1B). Immunoblot analysis showed that the Bs3-GFP and Bs3_S211A_-GFP fusion proteins were equally abundant *in planta* suggesting that the S211A mutation affects Bs3 function but not protein stability (Fig. 1C). Therefore, the serine to alanine mutation within the NADPH binding site resulted in a Bs3 derivative that does not trigger HR *in planta*, yet possible differences in NADPH oxidase activity of Bs3 and Bs3_S211A_ remained to be determined.

### Expression and purification of recombinant Bs3 and Bs3S211A protein

Plant immune reactions generally correlate with the release of H_2_O_2_ (Torres, 2010; Waszczak et al., 2018), however, our observation that Bs3-triggered HR correlates with release of H_2_O_2_ (Fig. 1B) does not explain if the detected ROS is produced by Bs3 itself, or by a potential downstream immune signaling component that Bs3 recruits to trigger HR. To clarify if Bs3 indeed has NADPH oxidase activity, we studied recombinantly-expressed *Bs3* by *in vitro* studies. *Bs3* and *Bs3*_*S211A*_ were expressed in *E. coli* and soluble Bs3 and Bs3_S211A_ protein was affinity purified with yields of two milligram per liter of culture (Fig. 2). Due to its homology to YUC proteins, we expected that Bs3 requires a bound FAD cofactor for enzymatic activity. Indeed, supplementation of the lysis buffer with FAD cofactor was necessary to obtain active Bs3 protein and to increase protein yield. In contrast to the purification protocol established for the *Arabidopsis thaliana* YUC6 protein, which uses extraction buffers containing 0.5 M sodium chloride (Dai et al., 2013), Bs3 purification works best under low sodium chloride conditions (< 100 mM). The purified Bs3 and Bs3_S211A_ proteins were bright yellow and the UV-vis spectra show characteristic peaks similar to FAD. Notably, we observed a slight shift of the local maxima of Bs3_S211A_ (at 368 nm and 453 nm) compared to the wild type Bs3 protein (373 nm and at 448 nm; Fig. 2B). This observation is consistent with the expected slight structural changes in Bs3_S211A_ as compared to wild-type Bs3.

**Figure 2:**
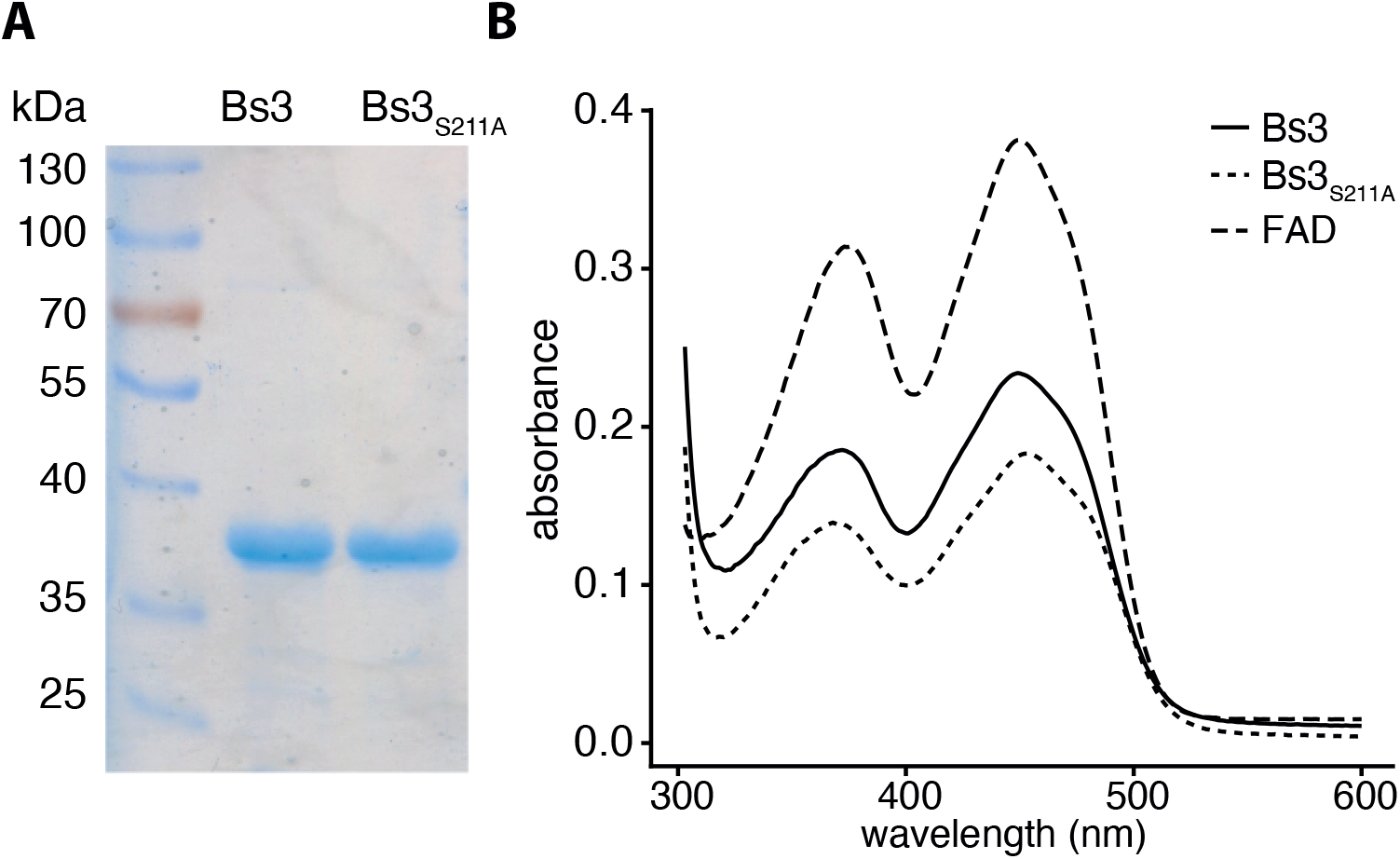
Recombinantly expressed Bs3 and Bs3_S211A_ bind the cofactor FAD. **A)** Recombinant Bs3-6xHis and Bs3_S211A_-6xHis protein was expressed in *E. coli*, purified using nickel columns, separated by size using SDS polyacrylamide gel electrophoresis and stained with Coomassie. B) UV-vis spectrum of FAD (dashed line), Bs3 (solid line) and Bs3_S211A_ (dotted line).

### Recombinant Bs3 and Bs3_S211A_ proteins oxidize NADPH and produce H_2_O_2_ in vitro

To compare NADPH oxidase activities of recombinant Bs3 protein and its mutant derivative Bs3_S211A_, we mixed corresponding protein fractions with the cofactor NADPH and monitored NADPH consumption via spectrophotometric measurements at 340 nm. The concentration of active protein was calculated from absorbance at 450 nm and the extinction coefficient of FAD (∊ = 11300 M^−1^ cm^−1^). At 25°C, we measured an NADPH oxidation activity of 63 nmol/mg*min for Bs3 and 137 nmol/mg*min for Bs3_S211A_ (Table 1, Fig. 3). These *in vitro* studies show that the Bs3_S211A_ mutant has approximately two-fold higher NADPH oxidase activity compared to that of the Bs3 wild type protein. This is in accordance with our expectation that the mutation of the conserved serine to alanine within the NADPH binding site destabilizes the C4a intermediate of Bs3_S211A_ and favors release of H_2_O_2_.

**Figure 3:**
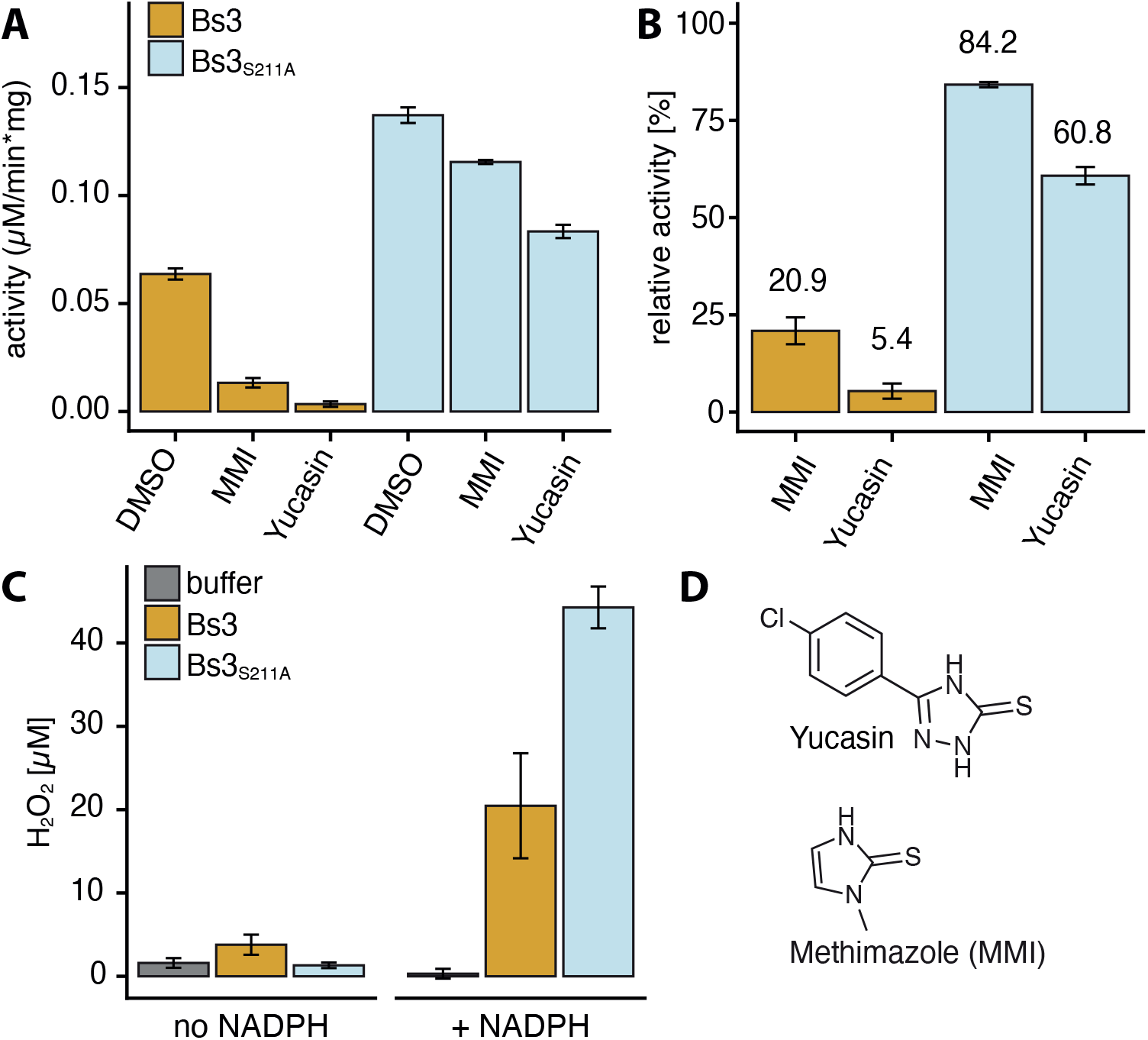
NADPH oxidation and H_2_O_2_ production by recombinantly-expressed Bs3 and Bs3_S211A_ protein. A) Buffer containing 100 μM NADPH was mixed with 0.2 μM Bs3 or Bs3_S211A_ and incubated at 25°C. NADPH oxidation was monitored via decrease of absorbance at 340 nm. Yucasin and methimazole (MMI) were dissolved in DMSO and added at a final concentration of 50 μM. B) Activity of Bs3 and Bs3_S211A_ was measured in presence of MMI or yucasin (from A). Measured values were normalized relative to the non-treated (DMSO) sample. C) 0.4 μM of depicted proteins was mixed with buffer containing no or 100 μM NADPH and incubated for five minutes at RT. The samples were subsequently mixed with HyPerBlu in a 1:1 ratio to measure H_2_O_2_ concentrations. Luminescence is measured after 10 min of incubation. Bars indicated mean +/s SD of three replicates. D) Chemical structures of yucasin and MMI.

**Table 1:** Yucasin and Methimazole (MMI) decrease NADPH oxidase activity of Bs3 and Bs3_S211A_

| Sample | Compound | Activity (nmol/mg*min) ± SD |
| --- | --- | --- |
| Bs3 | DMSO | 63.7 ± 2.6 |
| Bs3 | MMI | 13.3 ± 2.2 |
| Bs3 | YUCASIN | 3.4 ± 1.2 |
| Bs3<sub>S211A</sub> | DMSO | 137.2 ± 3.6 |
| Bs3<sub>S211A</sub> | MMI | 115.5 ± 0.9 |
| Bs3<sub>S211A</sub> | YUCASIN | 83.3 ± 3.1 |

### Competitive inhibitors of YUCs inhibit Bs3_S211A_ to a lesser extent than the Bs3 wildtype protein

We tested by *in vitro* assays if the two chemicals yucasin and methimazole (MMI), which are known as competitive inhibitors of YUC function (Nishimura et al., 2014), had an influence on NADPH oxidation by recombinant Bs3 or Bs3_S211A_ proteins. We observed that both Bs3 and Bs3_S211A_ have reduced NADPH oxidase activity upon inhibitor treatment (Fig.3). Yucasin and MMI reduced the NADPH oxidase of Bs3 activity by 95% and 81%, respectively (Fig. 3). These findings are in agreement with inhibitor studies on YUCs where yucasin was found to be a stronger inhibitor than MMI (Nishimura et al., 2014). Competitive inhibitors bind to the active site of the enzyme and prevent the substrate from binding. Thus, our observation that competitive inhibitors of YUCs also inhibit Bs3 suggests that the substrate binding sites of YUCs and Bs3 are to some degree structurally similar. It is worth noting that yucasin and MMI reduced the NADPH oxidase activity of the Bs3-derivative Bs3_S211A_ by only 29% and 16%, respectively (Fig. 3). The observation that both competitive inhibitors had less pronounced effects on Bs3_S211A_ as compared to the Bs3 wild-type protein possibly suggests that the S211A mutation affects the topology of the substrate binding site of Bs3.

### roGFP2 reporter assays indicate in planta NADPH oxidase activity for Bs3

The plant intracellular environment is likely distinct from the *in vitro* conditions in which we demonstrated NADPH oxidase activity for Bs3 (Fig. 3). To clarify whether or not Bs3 functions also *in planta* as an NADPH oxidase we utilized the reduction-oxidation-sensitive GFP-derivative roGFP2. Two cysteine residues in roGFP2 mediate its redox-sensitivity, which correlates with changes in its fluorescent properties ultimately allowing for ratiometric measurements in plant cells (Hanson et al., 2004; Schwarzlander et al., 2008). To measure predominantly Bs3-dependent redox changes we translationally fused roGFP2 to Bs3, thereby positioning the redox-reporter in spatial proximity of Bs3 (Fig. 4). The redox-reporter was also fused to the Bs3-derivative Bs3_S211A_ (mutation in NADPH binding site [G×G×**S**G ⟶ G×G×**A**G]; increased NADPH oxidase activity *in vitro*; see Fig. 3) and the previously studied Bs3 derivatives Bs3_G209A_ (mutation in NADPH binding site [Gx**G**xSG ⟶ Gx**A**xSG]) and Bs3_G41A_ (mutation in FAD binding site [G×**G**×SG ⟶ G×**A**×SG]; Krönauer et al 2019). Analysis of these distinct fusion proteins should clarify how distinct mutations in cofactor binding sites affect NADPH oxidase activity of Bs3 *in planta*.

**Figure 4:**
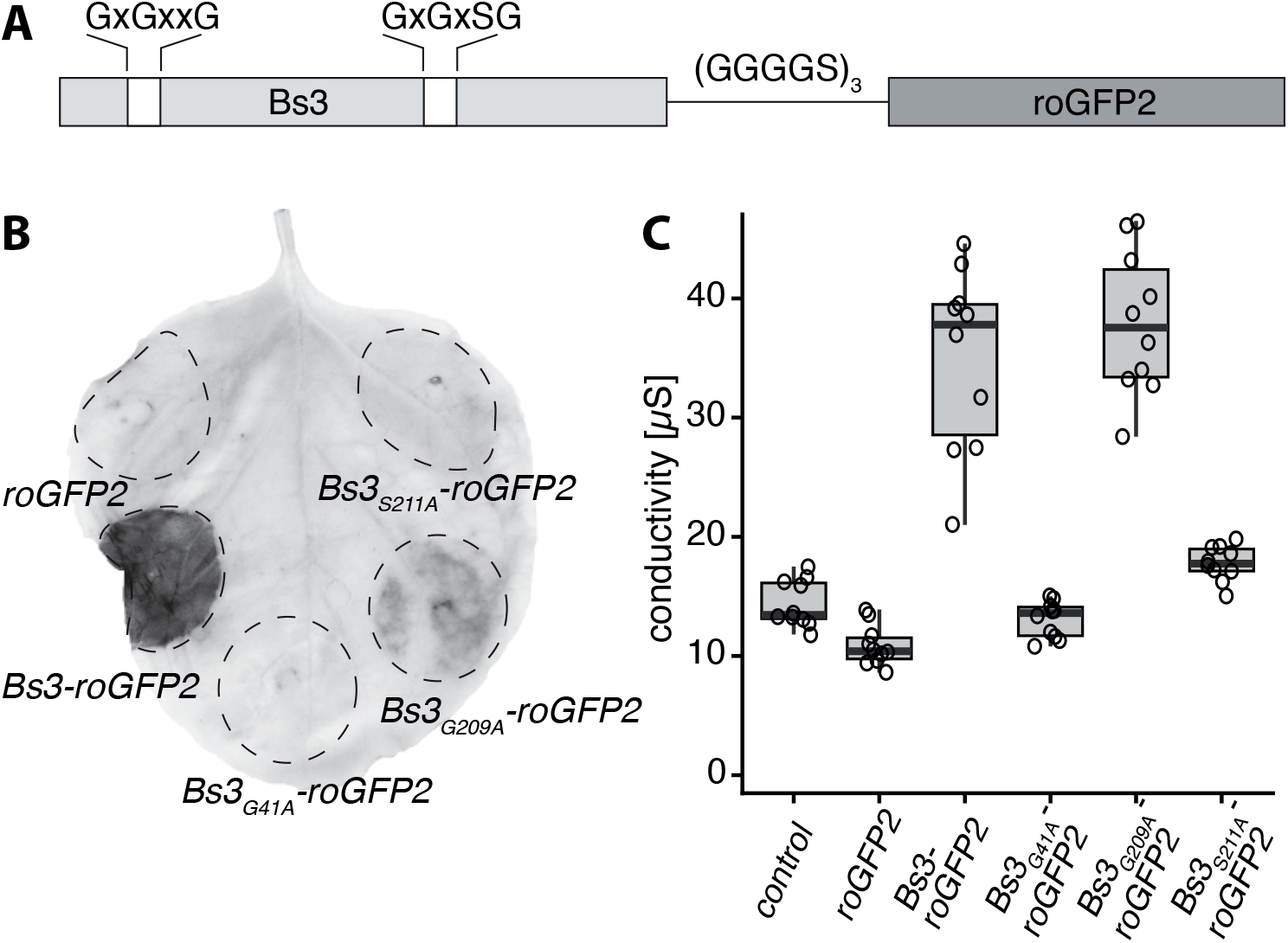
*35S* driven expression of *Bs3*_*G209A*_ triggers HR. A) Schematic representation of the Bs3-roGFP2 fusion construct. Gray Boxes on the left and right indicate the Bs3 protein and roGFP2 reporter, respectively. The connecting line indicates the linker with its amino acid sequence depicted above. Amino acid sequences above the white boxes indicate conserved sequences of FAD (left) and NADPH (right) binding sites of the Bs3 protein. B) *Agrobacterium* strains carrying the indicated constructs under control of the constitutive *35S* promoter were infiltrated into *N. benthamiana* leaves. Four days post infiltration, leaves were harvested and cleared with ethanol. Dashed lines mark the infiltrated area. Dark areas indicate presence of HR. C) Ion leakage measurements of plant tissue expressing *roGFP2* and *Bs3-roGFP2*. Depicted gene constructs that are under transcriptional control of the constitutive *35S* promoter were delivered into *N. benthamiana* leaves via *Agrobacterium*-mediated transient transformation. Three days post infiltration, leaf discs were harvested with a cork borer and incubated in ultrapure water. Conductivity was measured 20 hours post incubation. Boxplots represent measurements of 10 replicates. Each replicate is depicted as a black circle.

We first tested if the redox reporter roGFP2 had an impact on the functionality of translationally fused Bs3 protein and/or derivatives thereof by agroinfiltration of *35S* promoter-driven T-DNA constructs into *N. benthamiana* leaves. Two days post inoculation, leaves were cleared with ethanol to visualize the HR (Fig. 4). In addition, we carried out ion leakage assays to monitor HR. *35S*-promoter-driven *Bs3-roGFP2* induced visible cell death and high conductivity in *N. benthamiana* leaves, demonstrating that the roGFP2 reporter does not interfere with the Bs3-dependent HR (Fig. 4). Bs3_G209A_-roGFP2 caused similar electrical conductivity after three days but induced a less intense HR phenotype after four days compared to Bs3-roGFP2. The Bs3 cofactor-binding site mutants Bs3_G41A_-roGFP2 and Bs3_S211A_-roGFP2 did not induce HR nor elevated conductivity in agroinfiltration assays (Fig. 4). Overall, the observed HR phenotypes of the roGFP2 fusion proteins are consistent with the phenotypes observed for corresponding GFP fusion proteins (Fig. 1B, Krönauer 2019).

To analyze changes of the intracellular oxidation state caused by Bs3 and its mutant derivatives, the different *roGFP2* fusion constructs (*Bs3-roGFP2*, *Bs3_G41A_-roGFP2*, *Bs3_S211A_-roGFP2*, and *Bs3_G209A_-roGFP2*) and *roGFP2* were expressed in *N. benthamiana* leaves. Thirty hours post inoculation (hpi) confocal laser scanning microscopy (CLSM) was used to determine the ratios of fluorescence intensity (RFI) upon excitation at 405 and 488 nm (Fig. 5). We found that the HR-inducing proteins Bs3-roGFP2 and Bs3_G209A_-roGFP2 had about two-fold higher RFI values than the roGFP control (Fig. 5). This observation suggests that Bs3 and Bs3_G209A_ have indeed NADPH oxidase activity *in planta,* which would be consistent with a model where ROS produced by Bs3 triggers HR. Bs3_G41A_-roGFP2, which does not trigger HR, had similar RFI values as the roGFP2 control, still in agreement with our proposed model. However, the NADPH-binding site mutant Bs3_S211A_-roGFP2, which does not trigger HR *in planta*, had higher RFI values than the HR inducing Bs3-roGFP2 fusion protein. The conducted *in planta* NADPH oxidase studies of Bs3 and Bs3_S211A_ proteins are consistent with our *in vitro* studies that also showed elevated NADPH oxidase activity in Bs3_S211A_ as compared to the wildtype Bs3 protein (Fig. 3). In summary, we found that the mutant derivative Bs3_S211A_ does not trigger HR, but has higher NADPH oxidase activity as the Bs3 wild-type protein, which suggests that the release of H_2_O_2_ by Bs3 is not sufficient to trigger the Bs3-dependent HR.

**Figure 5.**
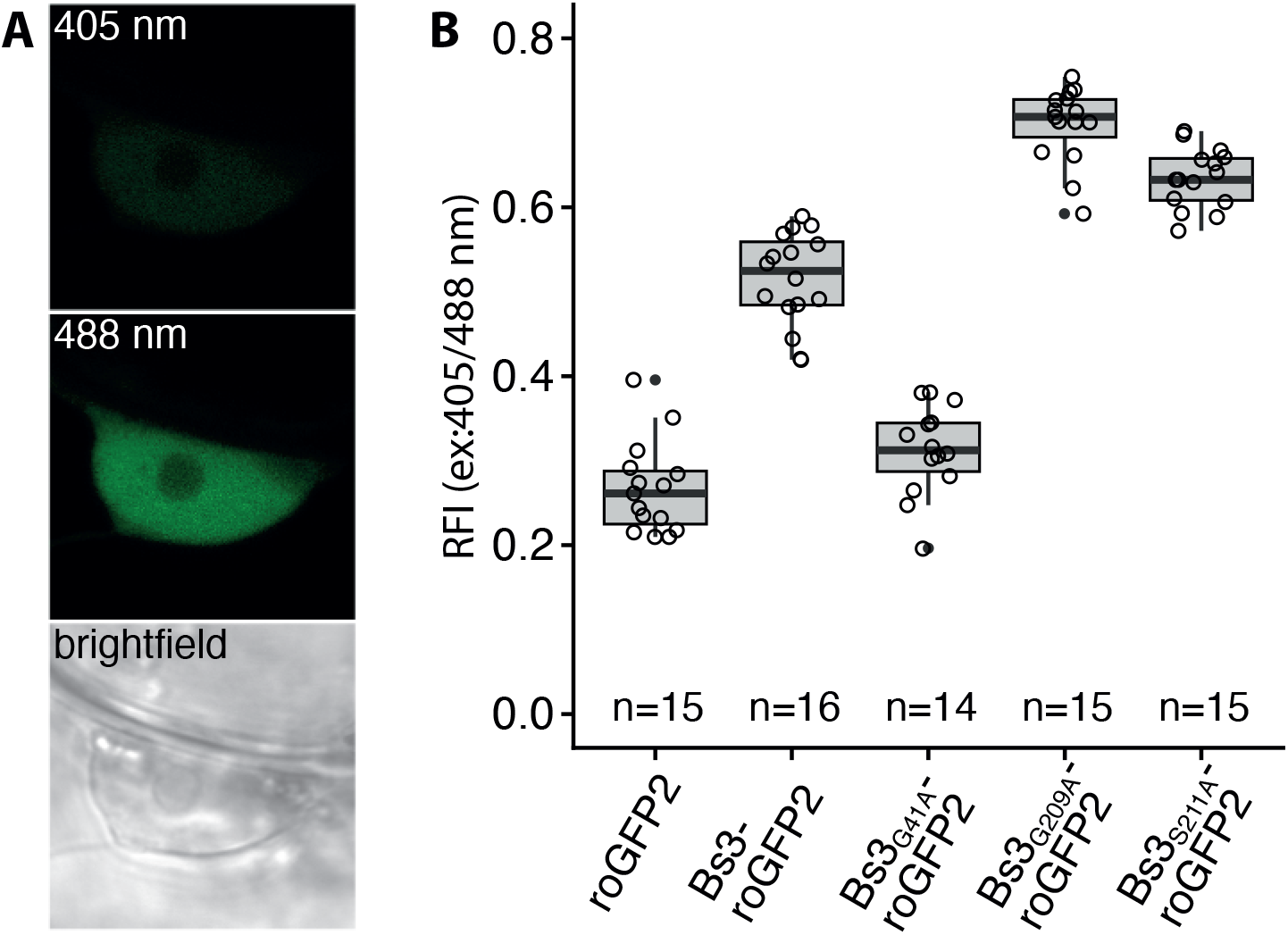
In planta oxidase activity of Bs3 and its derivatives. A) Representative pictures showing roGFP fluorescence in the nucleus with excitation at 405 and 488 nm. B) RoGFP2 oxidation is increased in *Bs3*, *Bs3*_*G209A*_ and *Bs3*_*S211A*_ expressing leaf tissue. Indicated gene constructs that are under transcriptional control of the constitutive *35S* promoter were delivered into *N. benthamiana* leaves via *Agrobacterium*-mediated transient transformation. 30 hpi leaf discs were analysed by ratiometric laser scanning microscopy and the ratios of fluorescence intensity (RFI) upon excitation at 405 and 488 nm was determined. n = number of measurements. Values corresponding to individual measurements are depicted as black circles.

### *Bs3* derivatives that trigger HR *in planta* cause growth arrest in yeast

Previous studies on executor-type R protein Xa10 from rice uncovered that Xa10 triggers cell death not only in plant cells but also in human cells (Tian et al., 2014). Inspired by this finding we wondered whether or not the executor R protein Bs3 would be functional in the budding yeast *Saccharomyces cerevisiae*, a model system of eukaryotic genetics. To study Bs3 function in yeast, we cloned the wild-type *Bs3* gene and mutant derivatives (*Bs3*_*S211A*_, *Bs3*_*G41A*_ and *Bs3*_*G209A*_) into a yeast expression vector under control of a galactose inducible promoter (*pGAL1*). Yeast transformants were grown in liquid medium, dropped onto either glucose (repressing) or galactose (inducing) containing agar plates and incubated at 30°C for 24 hours. We found that yeast containing wild-type *Bs3* grew only on glucose-but not on galactose-containing agar plates, suggesting that expression of wild-type *Bs3* inhibits proliferation of yeast cells (Fig. 6A). Growth reduction observed for yeast transformants expressing *Bs3*_*G209A*_, an HR-inducing *Bs3*-derivative, was nearly as pronounced as for strains expressing wild-type *Bs3*. By contrast, growth of strains expressing the *Bs3* derivatives *Bs3*_*S211A*_ or *Bs3*_*G41A*_, both of which do not trigger HR *in planta*, was similar to the yeast strain transformed with empty vector (EV) control.

**Figure 6:**
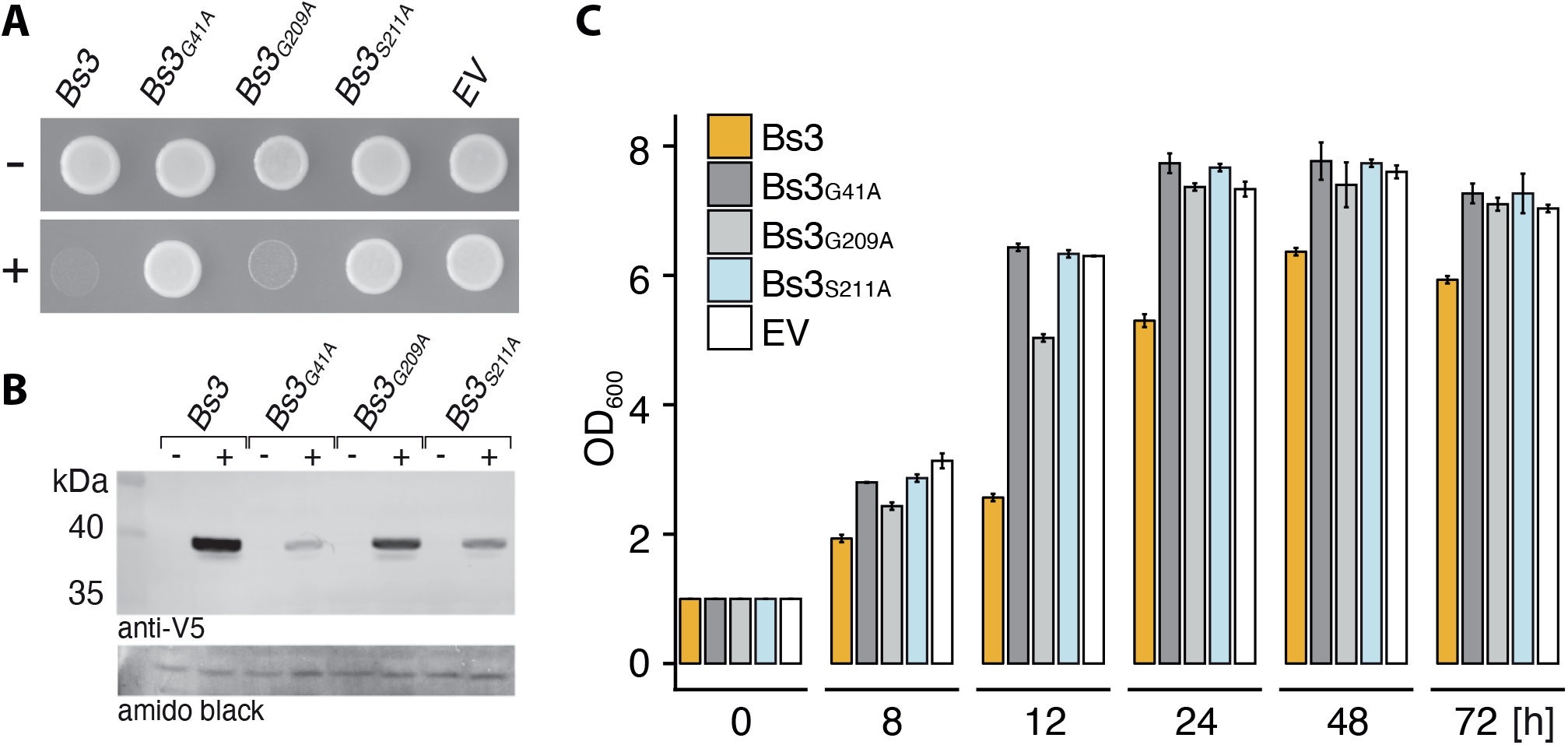
Bs3 but not Bs3_S211A_ impairs growth in yeast. Bs3 and *Bs3* mutant derivatives that are under transcriptional control of a galactose inducible promoter (pGAL1) were transformed into yeast. Subsequently, corresponding cultures were diluted to OD_600_ = 1, dropped onto repressing (−) or inducing (+) medium and incubated at 28°C for one day. A) Yeast cultures carrying the indicated gene constructs were grown in repressing (−) or inducing (+) liquid medium for 6 hours. Expression of depicted proteins was monitored via an anti-V5 immunoblot. Amido black staining was used to visualize total protein load. C) Yeast strains, carrying the indicated constructs, were grown in repressing medium overnight, diluted to OD_600_ = 1 in inducing medium and incubated at 28°C with shaking. Samples were taken at indicated timepoints and OD_600_ was measured. Values represent mean +/− SD of three replicates.

We also studied the effect of Bs3 and mutant derivatives on yeast grown in liquid medium. To do so, yeast cultures were diluted to a starting OD_600_ of 1 in inducing medium containing galactose. Expression of *Bs3* and its derivatives in corresponding yeast transformants were studied by immunoblot analysis (Fig. 6B) and revealed that Bs3 and Bs3_G209A_ protein levels were similar, while the mutant derivatives Bs3_G41A_ and Bs3_S211A_ showed somewhat lower abundance. Yeast growth in inducing liquid medium was monitored in a time-course experiment over a period of three days (Fig. 6C). In contrast to growth studies on agar plates, growth of yeast strains expressing the wild-type Bs3 protein was clearly detectable in liquid medium. However, consistent with the observed growth phenotypes on agar plates, yeast containing *Bs3* or *Bs3*_*G209A*_ showed reduced growth at 12 hours compared to the empty vector control strain. The yeast growth inhibition effect of Bs3_G209A_ was less severe than that of the wild-type Bs3 protein, which is in agreement with our previous observation that Bs3_G209A_ induced a less severe HR *in planta* than the wild-type Bs3 protein (Fig. 6C). Strains containing the mutant derivatives Bs3_G41A_ and Bs3_S211A_, that both do not trigger HR *in planta*, showed no yeast growth inhibition. In summary, Bs3 derivatives that trigger HR *in planta* also inhibit growth in yeast, suggesting that yeast provides a suitable platform to genetically dissect the molecular basis of Bs3-triggered cell death in future studies.

## DISCUSSION

### Competitive inhibitors of YUCCA proteins inhibit Bs3

Pepper Bs3 is one out of five currently known executor R proteins (Zhang 2015). Bs3 is exceptional since it is the only executor that shares sequence homology to proteins of known function. Due to the sequence homology of Bs3 to YUCCA proteins, which catalyze conversion of IPA to IAA, it would be possible that the enzymatic capabilities of Bs3 are similar to YUCCAs. Previously, we showed that Bs3, in contrast to the related YUCCA proteins, does not synthesize IAA and ultimately that Bs3-dependent HR does not involve changes in IAA levels (Krönauer 2019). It is still unanswered as to how the enzymatic features of YUCCA proteins are retained and/or different in Bs3 and could possibly explain why the expression of Bs3 triggers HR. Unfortunately, IPA, the metabolic substrate that YUCCAs convert into IAA, is a rather unstable metabolite that spontaneously converts into IAA *in vitro* (Gelinas-Marion et al., 2020). In consequence, quantification of putative IPA consumption by recombinant Bs3 protein in order to test if Bs3 and YUCCA use both IPA as a metabolic substrate *in vitro,* proves to be rather challenging.

Herein, we studied yucasin and methimazole, which are both competitive inhibitors of YUCCA proteins (Nishimura 2014). Given that both yucasin and methimazole reduced NADPH oxidase activity of both Bs3 and Bs3_S211A_ (Fig. 3), we assume that Bs3 has retained a substrate binding site that is structurally related to that of YUCCAs. The less severe inhibition of Bs3_S211A_ compared to Bs3 by yucasin and methimazole is in line with the notion that the S to A mutation does change the substrate-binding site but still allows NADPH oxidation.

### NADPH oxidase activity of Bs3 is not sufficient to trigger HR

Based on the fact that YUCCAs and other FMOs can synthesize H_2_O_2_ due to their inherent NADPH oxidase activity (Siddens et al., 2014; Thodberg et al., 2020), we hypothesized that Bs3 would produce larger quantities of H_2_O_2_ resulting in HR. We envisioned two possible ways in which Bs3 could trigger HR by the production of H_2_O_2_. Firstly, Bs3 might produce excessive amounts of H_2_O_2_ that cause oxidative damage to the plant cell, or alternatively, Bs3 produces H_2_O_2_ as a signaling molecule that activates a cascade leading to plant defense and HR. Indeed, our *in vitro* studies revealed that recombinant Bs3 protein produces substantial amounts of H_2_O_2_ (Fig. 3), and analysis of *in planta* expressed *Bs3* by roGFP2-based reporter assays (Fig. 4) indicate that Bs3 acts not only *in vitro* but also *in vivo* as an H_2_O_2_-producing NADPH oxidase.

To reinforce the possible causal link between direct production of H_2_O_2_ and HR we studied the function of Bs3 mutant derivatives. For example, the Bs3 mutant derivative Bs3_G41A_, which based on roGFP2 reporter assays, does not show detectable NADPH oxidase activity (Fig. 3) did also not trigger HR *in planta* (Fig. 4). This would be consistent with a model in which H_2_O_2_ produced by Bs3 triggers HR. To further support this hypothesis, we replicated a mutation in the fungal FMO SidA that was previously shown to cause increased NADPH oxidase activity (Shirey 2013). Indeed, *in vitro* and *in vivo* studies of the Bs3_S211A_ mutant showed increased NADPH oxidase activity (Fig. 3; Fig. 5). Despite the fact that Bs3_S211A_ has higher NADPH oxidase activity than the wild-type Bs3 protein, it does not trigger HR *in planta* (Fig. 4B). This implies that release of H_2_O_2_ by Bs3 is not sufficient to trigger HR. All in all, our current data suggests that Bs3 converts a metabolite which is possibly structurally related to IPA into a yet to be identified product that triggers HR. Whether or not this metabolic product of Bs3 is simply cytotoxic or if the metabolic product acts as a defense signaling molecule remains to be seen.

### Differentiation of direct and indirect ROS production during Bs3 HR

We studied Bs3-dependent H_2_O_2_ synthesis *in planta* via a roGFP2 reporter and by DAB staining. This raises the question whether or not the two systems detect the same or distinct H_2_O_2_ pools. Upon agroinfiltration of *35S*-*Bs3*, accumulation of Bs3 protein was detectable in *N*. *benthamiana* leaves at 24 hours post inoculation (hpi) and Bs3 levels peaked at ~36 hpi (Krönauer et al., 2019). RoGFP2 based measurements of the intracellular oxidation state were carried out at 30 hpi (Fig. 5). Notably, at this time, no DAB staining was visible (Fig. 5, Fig. S1), despite the fact that the fluorophore reporter suggested an increase in H_2_O_2_ at that timepoint. The intensity of DAB staining increased until the onset of cell death, even though a decrease of Bs3 protein levels was observed in *N. benthamiana* at 48 hpi (Fig. S1, Krönauer et al., 2019). It can therefore be assumed that the H_2_O_2_ that was detected by DAB staining was not produced by Bs3, but originated from other sources like membrane-bound NADPH oxidases or apoplastic peroxidases. In contrast to the DAB staining which is used to study plant tissue undergoing HR, typically correlating with strong H_2_O_2_ accumulation, our roGFP2 based measurements were able to detect changes in the oxidation state in parallel to Bs3-roGFP2 protein accumulation. The higher oxidation values measured for Bs3-roGFP2 compared to roGFP2 suggests that low amounts of H_2_O_2_ are produced directly by Bs3. Even though the higher oxidation values obtained for Bs3_S211A_-roGFP2 show that these low amounts of H_2_O_2_ are not sufficient to trigger cell death, it cannot be excluded that local changes in the oxidation state might contribute to the signaling cascade that subsequently triggers execution of HR.

### Bs3 derivatives that trigger HR in planta consistently inhibit proliferation in yeast cells

We found that expression of Bs3 limits proliferation of yeast cells, which are particularly amenable to genetic screens and can be used to dissect the Bs3-triggered cell death reaction in the future. The functional analysis of Bs3 and derivatives thereof in the plant- and yeast systems were consistent in all cases. For example, Bs3 and the NADPH-mutant derivative Bs3_G209A_ triggered HR *in planta* (Fig. 4) and also inhibited proliferation of yeast cells (Fig. 6). Bs3_G209A_ induced a somewhat weaker HR than the wildtype Bs3 protein *in planta* (Fig. 4B) and the same trend was observed in the yeast growth assay where expression of Bs3_G209A_ or Bs3 induced moderate and strong inhibition of yeast growth, respectively (Fig. 6A). Moreover, the Bs3 derivatives Bs3_G41A_ and Bs3_S211A_, did not trigger HR *in planta* and also failed to inhibit proliferation of yeast cells. The observed consistency of the Bs3-dependent phenotypes in yeast and plant cells possibly suggest that both phenotypes have a common molecular basis. Therefore, it stands to reason that the observed phenotypes in yeast and plant cell are caused by the same metabolite that Bs3 presumably produces in plant and yeast cells.

### Recombinant Bs3 protein as a tool to uncover the metabolic basis of the Bs3-triggered HR

Previously conducted biochemical studies uncovered that recombinant Arabidopsis YUCCA6 protein uses NADPH and oxygen to convert IPA to IAA (Dai 2013). Here, we established a protocol for purification of enzymatically active Bs3 protein that now enables us to compare enzymatic activity of Bs3 and the YUCCA proteins by *in vitro* assays. Given their high structural relatedness, it seems likely that Bs3 and YUCCA proteins are also similar with respect to their enzymatic features. Indeed, our studies revealed that Bs3, just like YUCCA6, has NADPH oxidase activity (Fig. 3A). Moreover, inhibitor studies suggest that YUCCA proteins and Bs3 have the same or at least structurally related metabolic substrates (Fig. 3). We envision future studies where incubation of metabolic candidate substrates, with enzymatically active Bs3 protein could provide the possibility to identify metabolic products of Bs3 by mass spectrometry. In summary, the established protocol for purification of recombinant enzymatically active Bs3 protein is the first step towards exploitation of its biochemical properties and holds the key to uncover the basis of Bs3-triggered immune reactions by metabolic studies.

## MATERIALS AND METHODS

### Plants and growth conditions

*N. benthamiana* and *Capsicum annuum* (cultivar ECW123) plants were grown at 20-24°C at 35-60% humidity with a light intensity of 12,3 klx and 16 hours light/ 8 hours dark cycle. Four to six weeks old plants were used for experiments.

### Plasmid construction

For *in planta* expression, the *Bs3* CDS was assembled with a *35S* promoter and *eGFP* into the *LIIa* expression vector via Golden Gate cloning (Binder et al., 2014). The serine to alanine mutation was introduced by PCR (PrimerFW: P-GCCGGGATCGATATCTCACTTG PrimerREV: ATTGCCACAGCCAACCGC). For bacterial expression, the Bs3 coding sequence was cloned into the pET-53-DEST (Novagen) expression vector via Gateway cloning. Protein solubility could be dramatically improved by deletion of the nucleotides encoding for the recombination site and the codons accounting for the two N-terminal methionines of Bs3. Deletion of these nucleotides was done by PCR mutagenesis (PrimerFW: P-GTGATGGTGGTGGTGATGTG and PrimerREV: AATCAGAATTGCTTTAATTCTTGTT CAC). For expression, in yeast the Bs3 CDS was cloned into the pYES-DEST-52 vector via Gateway cloning without further modification.

### DAB staining

Leaves are vacuum infiltrated with DAB staining solution (10 mM Sodium phosphate, 1 mg/ml Diaminobenzidine-tetrahydrochloride, 0.1% Tween-20, pH=7.2), incubated at room temperature with gentle shaking for at least 5 hours and subsequently de-stained with 80% ethanol at 60°C.

### *Xanthomonas* and Agrobacterium infiltration

*Xanthomonas* (82-8 uns*) carrying the pDSK602 vector with either *AvrBs3* or *AvrBs2* were grown at 28°C in NYG medium (5 g/L peptone, 3 g/L yeast extract, 20 g/L glycerol) containing Rifampicin and Spectinomycin at a final concentration of 100 μg/ml for 1 day. Cultures were pelleted and resuspended in water to OD_600_ = 0.4. Leaves of *C. annuum* were infiltrated with a blunt end syringe from the abaxial side.

*Agrobacterium tumefaciens* (GV3101) carrying the respective binary plasmids were grown overnight at 28°C in YEB medium (5 g/L beef extract, 1 g/L yeast extract, 5 g/L peptone, 5 g/L sucrose, and 0.5 g/L mM MgSO_4_, pH 7.2) containing Rifampicin and Spectinomycin at a final concentration of 100 μg/ml. Cultures were pelleted and resuspended in water to OD_600_ = 0.4. Leaves of *N. benthamiana* were infiltrated with a blunt end syringe from the abaxial side.

### Protein expression

*E. coli* Rosetta (Novagen) were transformed with *pET-53-DEST_Bs3* or derivatives, plated on LB Agar containing Ampicillin (100 μg/ml) and Chloramphenicol (15 μg/ml) and incubated at 37°C overnight. Several colonies were pooled to inoculate LB medium (10 g/L NaCl, 5 g/L yeast extract, 10 g/L tryptone) supplemented with Ampicillin (100 μg/ml) and Chloramphenicol (15 μg/ml). After overnight incubation at 37°C and 180 rpm, this starter culture was used to inoculate 2 L TB Medium (24 g/L yeast extract, 20 g/L tryptone, 4 ml/L glycerol, 0.072 M K_2_HPO_4_, 0.017 M KH_2_PO_4_) supplemented with Ampicillin (100 μg/ml) at a starting OD_600_ of 0.05. The culture was incubated at 37°C with shaking at 120 rpm until it reached an OD_600_ of 1. Cultures were then cooled down for 15 min in ice water and protein expression was induced with a final concentration of 1 mM Isopropyl 1-thio-β-D-galactopyranoside (IPTG). The cells were incubated for another 2,5 h at 18°C with shaking and followed by centrifugation (4500 x g, 30 min, 4°C). Pellets were stored at −20°C until further use.

### Protein purification

The bacterial pellet was re-suspended in lysis buffer (50 mM Potassium phosphate, 10% glycerol, 30 mM Imidazole, 1% Tween, protease inhibitor, 1 μM FAD, 1 mM DTT, pH = 8) using 5 ml of buffer per gram of pellet. 30 ml cell suspension were sonicated for 5 min (5s on/10s off, 60% amplitude) with a sonicator (EpiShear, Active Motif) equipped with a ¼” microtip probe. The lysate was centrifuged (16000xg, 4°C) for 30 min to pellet cell debris. An ÄKTA Pure 25 FPLC system, equipped with a 5 ml HisTrapFF Crude Column (GE Healthcare), was used for affinity purification. After column equilibration with 10 column volumes (CV) of wash buffer (50 mM Potassium phosphate, 10% Glycerol, 30 mM Imidazole, pH=8), the supernatant was loaded onto the column and washed with 20 CV wash buffer. The protein was eluted with 2 CV elution buffer (50 mM Potassium phosphate, 10% Glycerol, 500 mM Imidazole, pH=8). Imidazole was removed by dialysis and the protein was frozen in liquid nitrogen and stored at - 80°C until further use.

### NADPH oxidation

Spectroscopic assays were carried out in quartz cuvettes with a UV-900 UV-vis spectrometer (Shimadzu) equipped with a temperature controlled cell holder (TCC-100) set to 25 °C. Samples were diluted with 50 mM potassium phosphate buffer (pH = 8) containing NADPH. Exact NADPH concentrations were calculated from its absorption and the extinction coefficient at 340 nm (∊_340_ = 6220 M^−1^ cm^−1^).

### Detection of H_2_O_2_ with HyPerBlu

Protein was mixed with NADPH solution (100 μM) and incubated at RT for 15 min. Per replicate, 5 μl of this solution were transferred to a white 385 well plate, mixed with 5 μl of HyPerBlu solution (Lumigen) and incubated in darkness for 15 min at RT. Subsequently, luminescence was measured using a Berthold Tristar LB 941 plate reader. H_2_O_2_ concentrations were calculated using a standard curve prepared with known concentrations of H_2_O_2_.

### Redox Reporter Microscopy

Constructs in which *roGFP2* (Hanson et al., 2004) was fused to *Bs3* and its derivatives were transiently expressed in four to six week old *N. benthamiana* plants via *Agrobacterium* mediated transient transformation. 30 hpi, images were acquired using a Leica TCS SP8 confocal microscope by successive excitation at 405 nm and 488 nm and emission at 498 to 548nm. Images of nuclei were taken using a 63x water immersed objective and 10x digital magnification. Argon laser intensity was adjusted in a way so that pixels were close to saturation in samples with highest *roGFP2* expression. UV laser intensity was adjusted allowing imaging of samples with lowest *roGFP2* expression. Fiji (Schindelin et al., 2012) was used to crop the surrounding of the nuclei and to calculate mean pixel intensity of the fluorescent area.

### Ion leakage measurements

Ion leakage measurements were conducted using the CM100-2 conductivity meter (Reid & Associates). Each well was filled with 1 ml ultrapure water. Leaf discs (Ø 4 mm) were harvested three days post infiltration. One disc was added per well and incubated at RT. Ion leakage was measured after 20 hours of incubation.

## ACKNOWLEDGEMENTS

We thank D. Holmes for helpful comments on earlier versions of the article

## Notes

**Funding information** This work was supported by the Deutsche Forschungsgemeinschaft (SFB 1101 project D08 to T.L. and DFG grant no. LA1338/10-1).

### Competing Interest Statement

The authors have declared no competing interest.

